# Montane species track rising temperatures better in the tropics than in the temperate zone

**DOI:** 10.1101/2020.05.18.102848

**Authors:** Benjamin G. Freeman, Yiluan Song, Kenneth J Feeley, Kai Zhu

## Abstract

Many species are responding to global warming by shifting their distributions upslope to higher elevations, but the observed rates of shifts vary considerably among studies. Here we test the hypothesis that this variation is in part explained by latitude, with tropical species being generally more responsive to warming temperatures than are temperate species. We find support for this hypothesis in each of two independent empirical datasets—shifts in species’ elevational ranges, and changes in composition of forest inventory tree plots. Tropical species are tracking rising temperatures 2.1–2.4 times (range shift dataset) and 10 times (tree plot dataset) better than their temperate counterparts. Models predict that for a 100 m upslope shift in temperature isotherm, species at the equator have shifted their elevational ranges 93–96 m upslope, while species at 45° latitude have shifted only 37–42 m upslope. For tree plots, models predict that a 1°C increase in temperature leads to an increase in community temperature index (CTI), a metric of the average temperature optima of tree species within a plot, of 0.56 °C at the equator but no change in CTI at 45° latitude (–0.033). This latitudinal gradient in temperature tracking suggests that tropical montane biotas may be on an “escalator to extinction” as global temperatures continue to rise.

## Introduction

One consequence of global warming is the upslope shift, or “migration”, of many montane species’ ranges to higher, cooler elevations (Parmesan & Yohe 2003; Chen *et al*. 2011). However, there is significant variation in the degree to which montane species are effectively tracking temperature increases via distributional shifts to higher elevations. For example, while moths in Borneo are shifting their ranges upslope at rates that approximately match local warming rates (Chen *et al*. 2009; Wu *et al*. 2019), the upslope shifts in the ranges of plants in the European Alps lag far behind the pace of warming (Rumpf *et al*. 2018), and some birds in northeastern North America are shifting their ranges *downslope* despite recent warming (Zuckerberg *et al*. 2009; DeLuca & King 2017).

We address the question “why are some species effectively tracking changes in local temperatures along mountain slopes while others are not?” Observed elevational range changes associated with warming reflect a complicated set of factors including rates of warming, multidimensional changes in local climate (e.g., interacting effects of different climate variables and microclimate changes; Tingley *et al*. 2012; Zellweger *et al*. 2020), species’ evolutionary ecologies (e.g. functional traits; Angert *et al*. 2011; MacLean & Beissinger 2017), land use changes (e.g., anthropogenic and natural disturbances; Guo *et al*. 2018), stochastic events, and study methodologies (e.g., sampling effort and measurement error; Zhu *et al*. 2014). Here we focus on the possibility that biogeography shapes geographic response to recent warming; specifically, we test the hypothesis that observed variation in temperature tracking is explained in part by latitudinal position, with tighter temperature tracking in the tropics (reviewed by Freeman & Class Freeman 2014; Sheldon 2019).

The hypothesis that tropical species track changes in temperature better than their temperate counterparts is based on evidence that tropical species generally are physiologically more sensitive to temperature than temperate-zone species. Tropical ectotherms tend to live closer to their optimal temperatures and exhibit narrower thermal tolerances than temperate species (Deutsch *et al*. 2008; Huey *et al*. 2009; Dillon *et al*. 2010; Sunday *et al*. 2012; Perez *et al*. 2016; Polato *et al*. 2018). This physiological specialization appears to limit species’ elevational distributions, which are narrower in the tropics for both ectotherms and endotherms (McCain 2009). Tropical ectotherms’ thermal tolerances correspond more closely with the temperature conditions they experience within their elevational ranges compared to temperate-zone species, suggesting greater local adaptation to temperature in the tropics, where temperature seasonality is minimal (García-Robledo *et al*. 2016; Polato *et al*. 2018). These observations are consistent with a stronger role for temperature in controlling species’ elevational distributions in tropical mountains than in temperate mountains, leading to the expectation that tropical montane species will respond to warming temperatures by shifting their elevational ranges upslope more so than temperate montane species. Several studies have reported cases where tropical montane species are rapidly moving upslope associated with recent warming (reviewed by Freeman & Class Freeman 2014; Sheldon 2019). It is unknown, however, whether these scattered reports are generalizable across taxa and continents.

We used two independent datasets to test the hypothesis that temperature tracking in montane taxa is related to their latitudinal position. First, we compiled a dataset of resurveys that have reported elevational range shifts for communities and species associated with recent warming, hereafter the “range shift dataset” (Fig. 1a, Table S1, Datasets S1 and S2). We use the term “community” to refer to a set of species within a particular taxonomic group that lives within a particular montane region (e.g., a bird community or a bee community). Second, we analyzed inventory data from repeatedly surveyed forest plots located in montane areas across the Americas that have experienced significant warming over the past several decades, hereafter the “tree plot dataset” (Fig. 1b). These two datasets have complementary strengths for inferring latitudinal patterns in recent temperature tracking. Range shift studies have the advantage of explicitly measuring changes in elevational distributions for individual species, with data available for many taxonomic groups, but have the disadvantage that studies use different methods to study different taxa. In contrast, forest inventories use standardized methods and provide comprehensive information about the composition of tree communities at given locations, but generally cover shorter time scales, represent only trees, and are not globally distributed. The combination of two independent datasets, with complementary strengths, and large sample sizes within each dataset, allow us to rigorously test the hypothesis that there is a latitudinal gradient in temperature tracking.

**Figure 1.**
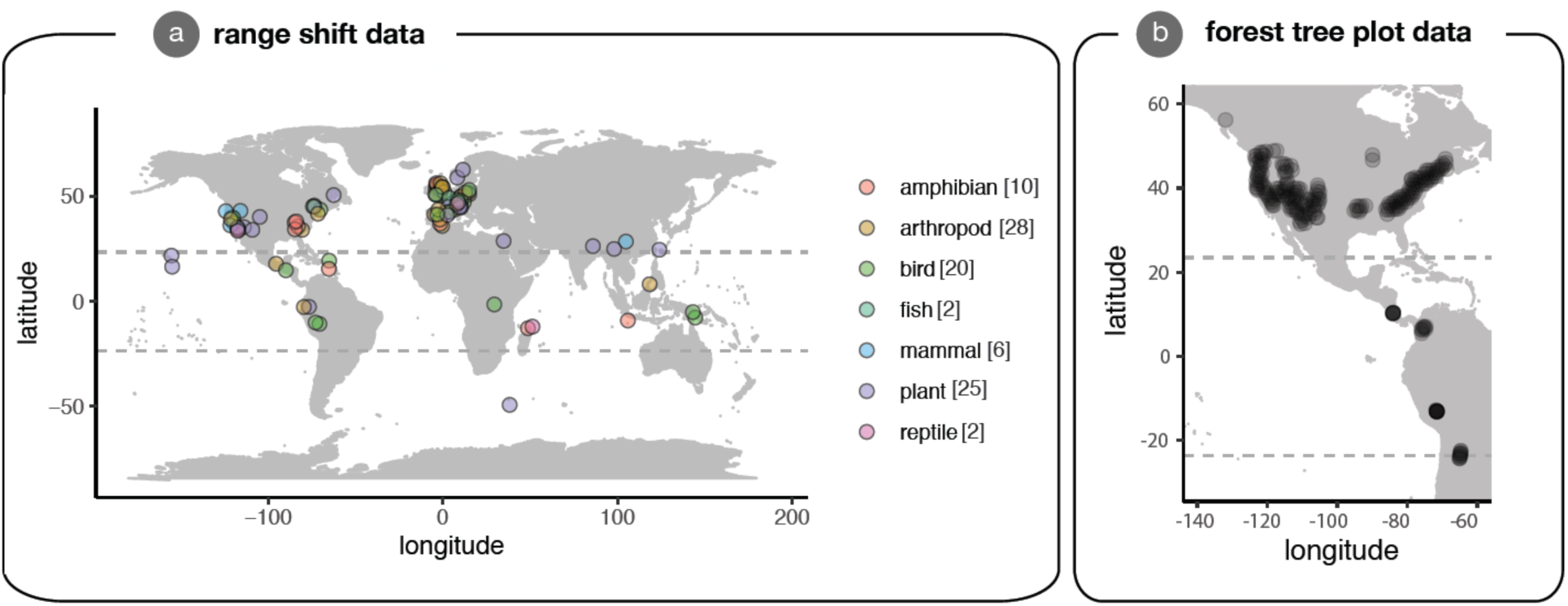
Maps of range shift studies that measured elevational shifts associated with recent warming (a) and of forest inventory tree plots that have been repeatedly censused (b). Locations of range shift studies are jittered slightly to improve clarity. The Tropics of Cancer and Capricorn (at 23.4**°** N and S, respectively) delimit the tropics, and are illustrated with dashed lines.

## Materials and methods

### Range shift dataset

We compiled a comprehensive list of studies that have measured elevational range shifts associated with recent warming (within the past ∼ 100 years). First, we conducted Web of Science searches on 11 July 2019 and 5 January 2021 with the keywords “climate change” OR “global warming” AND “range shift” AND “mountain” OR “elevation” OR “altitude*”. These searches returned 1827 and 2164 hits, respectively. We retained studies that met the following three criteria: (1) they measured recent range shifts at species’ lower elevation limits, mean (or optimum) elevations, or upper elevational limits; (2) range shifts were reported for all species or the entire community, not just species with significant range shifts; and (3) range shifts were measured over a time period of ≥10 years. Second, we located additional studies that met these three criteria by examining recent papers synthesizing the range shift literature (Chen *et al*. 2011; Lenoir & Svenning 2015; Wiens 2016; Freeman *et al*. 2018a; Rumpf *et al*. 2019; Lenoir *et al*. 2020). There were three cases for which multiple publications reported elevational range shifts for the same community using the same underlying data (plants in France, and butterflies and birds in Great Britain). For these cases, we included only the study with a larger sample size of species.

For each study that met our criteria (see Table S1, full data provided in Dataset S1), we extracted the following key information: (1) taxonomic group; (2) mean latitude of the study site; (3) duration over which range shifts were calculated (e.g., number of years elapsed between historic and modern surveys); (4) spatial scale of study (“local” when studies were conducted along single elevational gradients or entirely within small montane regions; “regional” when the study was conducted within large political units such as the state of California or the country of Spain); (5) number of species included in the study; (6) range shifts at lower elevational limits, mean/optimum elevations, and upper elevational limits for the entire community (i.e., for a community of 30 species, the mean range shift of these 30 species); (7) range shifts at lower elevational limits, mean/optimum elevations, and upper elevational limits for individual species (note that not all studies reported species-specific range shifts); (8) temperature changes at the study site between surveys; and, if reported, (9) expected elevational range shifts based on local temperature changes and lapse rate. There were two studies that did not report local temperature changes (Moret *et al*. 2016; Kusrini *et al*. 2017). For these studies, we estimated local temperature changes using gridded data provided by the Climatic Research Unit (Harris *et al*. 2013). For studies that did not report local adiabatic lapse rates, we used lapse rates reported for the geographically nearest study within our dataset, following Chen et al. (2011). When data were presented only in figures, we used WebPlotDigitizer to extract data from the published graphics (Rohatgi 2017).

### Range shift dataset: statistical analysis

Our primary analyses tested for latitudinal patterns in temperature tracking. We calculated the temperature tracking score as the ratio of the observed elevational shift to the expected elevational shift given estimates of local warming and adiabatic lapse rate. Temperature tracking scores of ∼1 indicate that observed changes closely match those expected based on concurrent warming (i.e., strong temperature tracking), and scores >1 indicate that species are shifting their ranges faster than expected. Conversely, temperature tracking scores that are positive but closer to 0 indicate upslope shifts in the ranges of species (or, for the tree plot dataset, the composition of tree plots) that lag behind expectations given rising temperatures. Negative scores indicate changes in species ranges or composition that are in the “wrong” direction (e.g., downslope shifts despite warming temperatures). Because we were interested in responses to warming, we did not include three range shift studies that reported cooling temperatures when analyzing temperature tracking (Moskwik 2014; Neate**-**Clegg *et al*. 2020). We analyzed the range shift data at both the community-level (data = 162 estimates of temperature tracking from 90 communities from taxonomic groups including plants, birds, insects, mammals and amphibians; Figure 1a, data provided in Dataset S1) and species-level (data = 6141 estimates of temperature tracking from 2951 species from 72 communities; species-level data was unavailable for 15 communities; see Figure S1, data provided in Dataset S2). All statistics were done in R version 3.6.2 (R Development Core Team 2020).

We analyzed latitudinal patterns in temperature tracking using both categorical and continuous approaches. We analyze latitude to follow the formulation of the hypothesis we are testing; note that latitude is a proxy variable for complex multidimensional variation in the underlying climatic and biotic variables that species directly experience (note that latitude and mean annual temperature, as well as mean annual temperature and other temperature metrics, are tightly correlated, see Figures S2 and S3). We first analyzed categorical differences between latitudinal zones, categorizing communities and species as being “tropical” or “temperate” based on the location of the study, with tropical locations defined as < 23.4° absolute latitude. We then fit linear mixed-effects models using the “lme4” package (Bates *et al*. 2014). The response variable in both community-level and species-level models was the temperature tracking score. We included latitudinal zone (tropical vs. temperate) and four methodological covariates as fixed effects: (1) distributional variable measured (lower limit vs. mean elevation vs. upper limit); (2) spatial scale of study (local vs. regional); (3) number of species in the study (for the community-level model only); and (4) duration of the study. We included the community ID (i.e., the study) as a random effect, as multiple distributional variables (i.e., changes at lower limits, mean elevations, and upper limits) were reported for many communities. For the species-level models, we included species name as a random effect. To investigate whether our community-level results were driven by the inclusion of communities with few species, we fit an additional model including data only from communities with ≥10 species (135 estimates of temperature tracking from 74 communities).

We analyzed latitudinal differences in temperature tracking as a continuous function of latitude by replacing the factor “tropical/temperate” with absolute latitude in models.

We did not include taxa as a predictor variable in any models because taxonomic differences predict minimal observed variation in recent range shifts (Chen *et al*. 2011; Lenoir *et al*. 2020). Latitudinal sampling was also poor for most taxonomic groups. For the one exception—birds—we repeated analyses after restricting our dataset to only bird studies (6 from tropics, 13 from the temperate zone). We also present patterns for the only other taxonomic groups with two or more studies from both tropical and temperate zones (amphibians, arthropods, and plants).

Last, we also quantified latitudinal patterns in absolute response to warming temperatures. We therefore repeated analyses with elevational shift (m/decade) as the response variable instead of temperature tracking.

### Tree plot dataset

We compiled our tree plot dataset using Forest Inventory and Analysis (FIA) plots from the United States and previously published inventory plot data from Central and South America (Feeley *et al*. 2013; Fadrique *et al*. 2018). We filtered for FIA plots that were fully forested and that have not received observable intervention, including cutting, site preparation, artificial regeneration, natural regeneration, and other silvicultural treatment (Smith 2002). We also filtered for FIA plots that have not experienced disturbances in at least five years. Because we were interested in changes in just mountain forests, we selected FIA plots that fell within mountainous areas using the global mountains raster map derived from the 250 m global Hammond landforms product (Karagulle *et al*. 2017). We included FIA plots that have been surveyed two or more times. We collected the species identity of all individual adult trees (diameter at breast height > 12.7 cm, or 5 inches) for each survey of each FIA plot. Our dataset of forest inventory plots contained 11,023 plots from temperate montane forests from the United States, and combined this FIA data with data from tropical montane forests from Central (10 sites in Costa Rica; Feeley *et al*. 2013) and South America (186 sites in the tropical Andes; Fadrique *et al*. 2018). The tropical tree plots were set up with similar criteria of no sign of recent interventions or disturbances, and were also located in mountainous areas. Only adult trees (diameter at breast height ≥ 10 cm) were surveyed in the tropical plots.

### Tree plot dataset: statistical analysis

For forest inventory data, we analyzed changes in plot-level community temperature indices (CTI) over time in relation to local warming. We calculated the CTI of the plot as the average of optimal temperature of each species weighted by basal area, following the method of Fadrique et al. (2018). Specifically, we downloaded all georeferenced plant location records available through the BIEN database (version 4.1.1 accessed in November 2018 via BIEN package in R) for the New World (North America, Central America, and South America, but excluding the Caribbean islands). The BIEN database provides collated observation and collection data from multiple sources and provides a base level of data filtering and standardization. We used BIEN’s default download preferences to exclude records of known introduced species and cultivated individuals. We further filtered the records to include only those that were georeferenced and that list the year of collection/observation as being between 1970 and 1980. We restricted records to just this 10-year window to minimize errors in quantifying temperature optima due to the possibility of species changing their ranges through time. For each species with ≥ 10 retained records, we then extracted the estimated mean annual temperature (BIOCLIM1) at all collection coordinates from the CHELSA v1.2 raster of “current” (i.e., mean of 1979-2012) climate at 30 arc-second resolution and estimated the species’ thermal optima (MAT_opt_) as the mean MAT. For species with < 10 records but >= 10 records from all congeners, we used the estimated MATopt at the genus-level. Species with < 10 records and < 10 records from congeners were excluded from the subsequent analyses. We additionally ran analyses using only species-level climatic optima measurements. We next used the collection records to calculate the CTI of each plot in each census as the mean MAT_opt_ for species in the plot weighted by their relative basal area.

We then calculated the thermophilization rate of each FIA plot as the linear trend of CTI over time, and combined this dataset with previously published thermophilization rates for tropical tree plots. For each plot, we calculated the rate of temperature change as the linear trend of mean annual temperature over time, using monthly mean temperature estimates from 1980 to 2013 from the CHELSA time-series dataset (Karger & Zimmermann 2019). As with the range shift data, we calculated the temperature tracking score for each plot as the ratio of observed changes (rate of change in thermophilization rate) and expected changes (rate of change of mean annual temperature).

In order to focus on the response of organisms to global warming, we selected plots with significant warming trends (plots with *p* < 0.05 in a regression of mean temperature versus year), leaving 8056 temperate plots and 44 tropical plots. As FIA plots are much smaller in size compared to the plots in Central and South America (673 m^2^ for FIA plots compared to 1-hectare for most tropical plots), we aggregated FIA plots into 1 degree diameter hexagons that contain approximately 27 plots each, averaging the thermophilization rate and temperature tracking score weighted by total basal area (Table S2). We removed hexagons with < 5 plots from subsequent analysis, leaving 212 aggregated temperate tree plots (Figures 1b and S4; data provided in Dataset S3). We examined the relationships between latitude and the temperature tracking score by fitting linear mixed-effects models using the “nlme” package in R (Pinheiro *et al*. 2017). We included both latitude (either the factor “tropical/temperate”, or absolute latitude) and elevation as fixed effects. Due to the close proximity of hexagons/plots and spatial dependence in the residuals of non-spatial linear models, we modeled spatial random effects that follow Gaussian covariance functions (Pinheiro *et al*. 2017), and evaluated the significance of regression coefficients for fixed effects using the conditional standard error of regression coefficients.

## Results

### Range shift dataset

Tropical montane taxa are tracking temperature increases “better” (i.e., have temperature tracking scores closer to 1) than are temperate montane species (Figure 2a-d, Fig. S5). Tropical communities had temperature tracking scores 2.4 times greater than temperate communities (Figure 2a, Table S3; temperature tracking scores for tropical and temperate communities = 0.85 ± 0.15 vs. 0.36 ± 0.070; df = 71.5, *t* = −2.99, *p* = 0.0039; estimates ± standard errors from mixed-effects models), and tropical species had temperature tracking scores 2.1 times greater than temperate species (Figure 2c, Table S4; temperature tracking scores for tropical and temperate species = 0.87 ± 0.18 vs. 0.41 ± 0.079; df = 64.9, *t* = −2.47, *p* = 0.016). Results were similar when we modeled temperature tracking as a continuous function of position along the latitudinal gradient. Temperature tracking scores decreased by an average of 0.13 ± 0.048 and 0.12 ± 0.053 per 10° increase in absolute latitude for communities and species, respectively (Figures 2b and 2d, Tables S5-S6). For range shift data, estimates from linear mixed models are that communities have a tracking score of 0.96 ± 0.20 at the equator but 0.37 ± 0.069 at 45° latitude (estimates ± standard errors, averaged over levels of fixed effects; the equivalent values for the species-level model are 0.93 ± 0.22 and 0.42 ± 0.082). Hence, tropical communities and species are closely tracking temperature changes while temperate zone communities and species are not, though the explanatory power of community models was much greater (marginal R^2^ values from linear mixed-effects models were 0.11 and 0.098 for categorical and continuous community models, respectively, versus and 0.0037 and 0.0091 for species-level models).

**Figure 2.**
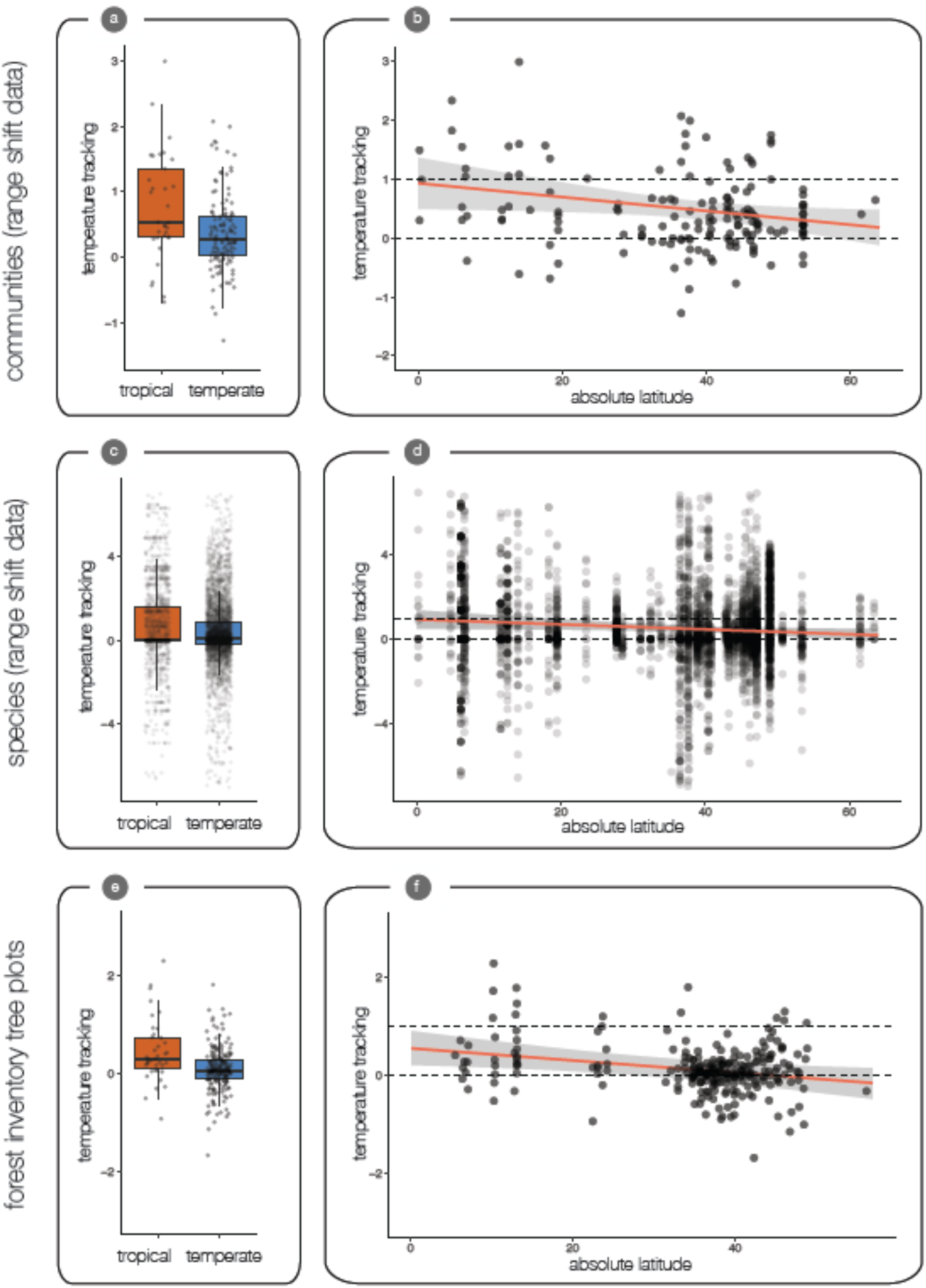
Tropical communities, species, and forest inventory tree plots have higher temperature tracking scores than their temperate zone counterparts. Raw data are shown as points. Dashed lines illustrate both perfect temperature tracking (temperature tracking = 1) and zero temperature tracking despite warming temperatures (temperature tracking = 0). Trendlines illustrate predictions from mixed models; shaded areas illustrate 95% prediction intervals. (a) temperature tracking for communities in tropical and temperate zones; (b) relationship between temperature tracking and absolute latitude (°) for communities; (c) temperature tracking for species in tropical and temperate zones; (d) relationship between temperature tracking and absolute latitude (°) for species (note that species with extreme temperature tracking values have been removed to improve visualization in panels c and d); (e) temperature tracking for forest inventory tree plots in tropical and temperate zones; (f) relationship between temperature tracking and absolute latitude (°) for forest inventory tree plots.

Methodological covariates included in models had minimal explanatory power, with the exception that temperature tracking scores at species’ upper elevational limits were higher compared to temperature tracking scores at their lower elevational limits or mean elevations (Tables S3-S6). Results for communities all held when considering only communities with 10 or more species (Figure S6, Tables S7-S8), indicating that the results are not driven by the inclusion of depauperate communities. All results held when subsetting the range shift dataset to only studies of birds (Figure S7, Tables S9-S12), and patterns were similar for other taxonomic groups with available data from both tropical and temperate zones (Fig. S8).

The magnitude of recent warming has been greater at high latitudes (IPCC 2014), and nearly all studies in our dataset that report fast rates of recent warming are from the temperate zone (Figure S9). However, greater absolute warming in the temperate zone did not lead to greater absolute upslope shifts (in units of m/ decade) in the temperate zone. Instead, due to tighter temperature tracking in the tropics, model estimates of absolute shifts (in units of m/ decade) were slightly greater in the tropics (24.7 ± 6.33 m/decade) than the temperate zone (14.7 ± 2.88 m/decade); this difference was not statistically significant (df = 77.8, *t* = -1.49, *p* = 0.14; Figure 3a-d, Tables S13-S16; estimated upslope shift from a model that did not include latitudinal zone was 25.13 ± 5.37 m/decade).

**Figure 3.**
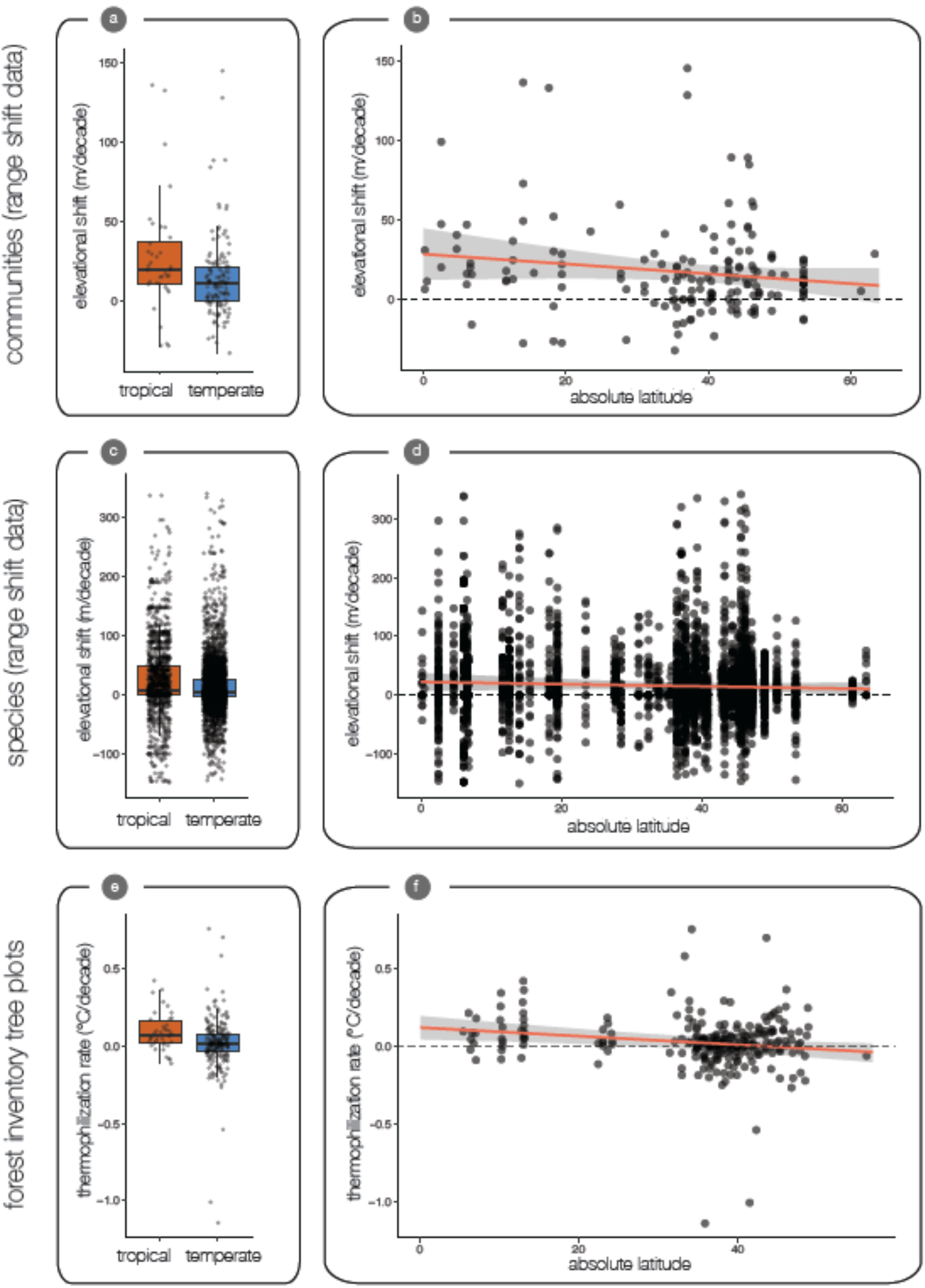
Tropical communities and species have undertaken larger upslope shifts, and forest inventory tree plots greater thermophilization, than their temperate zone counterparts. Raw data are shown as points. Trendlines illustrate predictions from mixed models; shaded areas illustrate 95% prediction intervals. (a) elevational shifts for communities in tropical and temperate zones; (b) relationship between elevational shifts and absolute latitude (°) for communities; (c) elevational shifts for species in tropical and temperate zones (note that species with extreme temperature tracking values have been removed to improve visualization); (d) relationship between elevational shifts and absolute latitude (°) for species (note that species with extreme elevational shift values have been removed to improve visualization in panels c and d); (e) thermophilization rate for forest inventory tree plots in tropical and temperate zones; (f) relationship between thermophilization rate and absolute latitude (°) for forest inventory tree plots.

### Forest inventory tree plot dataset

Tropical montane trees are also tracking temperature increases better than temperate montane trees (Figure 2e-f, Figure S10, Tables S17-S18). Tropical montane forest plots had higher temperature tracking scores (0.44 ± 0.11) than temperate montane forest plots (0.044 ± 0.044; df = 253, *t* = −3.42, *p* = 0.00072; Table S17). Latitude is a significant predictor of temperature tracking in the linear mixed model: a 10° increase in absolute latitude corresponds to a 0.13 ± 0.039 decrease in the temperature tracking score (Table S18). The model-based estimate is that tree plots have a tracking score of 0.56 ± 0.14 at the equator, but −0.033 ± 0.14 at 45° latitude. Results were unchanged when using only species-level climatic optima data (Tables S19-S20) or when using alternative methods for aggregating FIA plots (Tables S21-S24).

The greater temperature tracking in tropical inventory plots was due to faster changes in CTI, also known as the thermophilization rate, in the tropics. Tropical plots had thermophilization rates of 0.095 ± 0.020 °C per decade vs. 0.017 ± 0.012 °C per decade for temperate plots (df = 253, *t* = −2.74, *p* = 0.065; Figure 3e,f). Thus, while the rate of warming was faster at high latitudes in our tree plot dataset—0.31 ± 0.0065 °C per decade for temperate plots vs. 0.22 ± 0.010 °C per decade for tropical plots (df = 253, *t* = 3.19, *p* = 0.0016; Figure S11)— there were still greater absolute responses to recent warming in the tropics.

## Discussion

Biogeography predicts how montane species are changing their elevational ranges as temperatures rise. Species are on the move at low latitudes, where tropical species are, on average, closely tracking recent temperature increases by shifting their distributions upslope. In contrast, temperate species’ elevational ranges are shifting upslope at rates that lag far behind the pace of warming. These results, replicated in both range shift and tree plot datasets, provide evidence that species’ elevational distributions are more tightly associated with mean temperature in the tropics than the temperate zone, as has been previously hypothesized in other contexts (e.g., Ghalambor *et al*. 2006; Polato *et al*. 2018).

The similar results from independent range shift and tree plot datasets bolster our confidence that the latitudinal gradient in temperature tracking is real. Nevertheless, while both datasets had similar estimated slopes for the relationship between temperature tracking and latitude, the range shift dataset had a higher intercept. This difference could reflect a biological difference in generation time between species included in the datasets. Trees have long generation times that lead to slow rates of community turnover and range shifts (Lenoir *et al*. 2008; Feeley *et al*. 2012). In comparison, the range shift dataset consists primarily of taxa such as birds, mammals, insects and herbaceous plants that typically have shorter generation times than trees. An alternative explanation is that methodological differences between range shift studies and tree plots, both in data collection and analysis, explain why temperature tracking scores are higher in the range shift dataset.

The results of the range shift analysis appear to be robust to the heterogeneity present within this dataset. Despite nearly two decades of research documenting elevational range shifts associated with recent warming, the number of studies for most taxonomic groups remains low, particularly in the tropics (Feeley *et al*. 2017). Birds are the only taxonomic group with reasonable sampling across temperate and tropical zones, and we find strong temperature tracking in tropical—but not temperate—birds. This means that we have evidence for a latitudinal gradient in temperature tracking in both birds and trees (comparing tropical birds to temperate birds, and tropical trees to temperate trees). Patterns for other taxonomic groups (e.g., amphibians, arthropods, and plants) appear to be similar, but are provisional given the limited amount of data currently available from the tropics. We hope that our analysis motivates further research measuring how tropical montane species have responded to recent warming.

### What generates the latitudinal gradient in temperature tracking?

Multiple mechanisms may explain why tropical species are tracking temperature changes better than temperate species. The leading explanation is that tropical species are more physiologically sensitive to climate change than are temperate species (e.g., Deutsch *et al*. 2008). The purported heightened sensitivity of tropical species is supported by the fact that tropoical species generally inhabit distributions that experience narrower ranges of temperature, accounting for inter and intra-annual variation, than their temperate counterparts. An additional possibility is that both tropical and temperate species are tracking recent warming, but that temperate species are using phenological shifts to do so (Socolar *et al*. 2017). Seasonal temperature fluctuations in the tropics are minimal, meaning that tropical species are unlikely to be able to track climate via phenological shifts. In other words, temperate-zone species may track changing climate by shifting in time, while tropical species track changing climate by shifting in space. Last, an alternative explanation is that elevational specialists are particularly responsive to warming regardless of latitude, but that such species predominate in the tropics, where high species richness creates stronger interspecific competition (and other species interactions) that leads species to inhabit narrower elevational distributions. That is, mean annual temperature is a more important driver of species’ elevational distributions in the tropics, but acts through indirect mechanisms (e.g. species interactions that restrict elevational distributions). Consistent with this view, the lowest temperature tracking scores within the tropics in our dataset come from species-poor tropical islands such as Hawaii and Puerto Rico, while temperature tracking scores average higher on species-rich tropical mainlands.

### Variation in temperature tracking is substantial

We report strong statistical support for the existence of a latitudinal gradient in temperature tracking. Nevertheless, our ability to explain variation in observed temperature tracking remains limited, and is contingent on the scale of analysis. Latitude is a much better predictor of temperature tracking when considering communities (a set of species aggregated together) than for individual species, which show a wide variation in their temperature tracking scores (Freeman *et al*. 2018a; Rumpf *et al*. 2019). These results indicate that temperature is only one of many factors that drive species’ elevational range shifts; other potentially important factors include species interactions, measurement error, stochastic events, interacting effects of different climate variables, microclimate changes, land use changes, and disturbances. Consequently, despite a clear latitudinal pattern of temperature tracking for communities, we still have little ability to predict how individual species’ elevational distributions are changing associated with warming temperatures (Angert *et al*. 2011). Range shifts for individual species may be more predictable in the marine realm (Pinsky *et al*. 2019; Lenoir *et al*. 2020).

### Limitations

Several limitations of our study deserve explicit mention. First, we followed previous analyses in calculating temperature tracking scores based on mean annual temperature (Chen *et al*. 2011). Analyses that incorporate temperature variability (i.e., seasonality) and other climatic variables have also proven powerful (Crimmins *et al*. 2011; Tingley *et al*. 2012), but are inherently more difficult to implement and interpret. Second, our analyses do not take into account variation in microclimate, which may be a strong driver of range shifts or the lack thereof (Lembrechts *et al*. 2019; Zellweger *et al*. 2020). It is not clear how microclimate availability varies along a latitudinal gradient, but a greater availability of microclimates and climate refugia in the temperate zone than the tropics is an alternative explanation for our results. We were similarly unable to analyze climatic factors that occur at intermediate spatial scales along mountain slopes, such as cold-air pooling (Curtis *et al*. 2014). Third, future studies should address landscape-level changes due to habitat loss or other disturbances (Larsen 2012; Lenoir & Svenning 2015; Campos-Cerqueira *et al*. 2017; Guo *et al*. 2018). The range shift resurveys and forest plots in our dataset took place in landscapes that have not undergone intensive deforestation or other land-use change. Given that highly modified landscapes predominate across most of the globe, further tests of the interactions between landscape change and climate change are needed. Fourth, the patterns we document for communities are not without exceptions. For example, Puerto Rican frogs and birds are not closely tracking temperature despite their tropical latitude (Campos-Cerqueira & Mitchell Aide 2017; Campos-Cerqueira *et al*. 2017), though this could potentially reflect increasing forest cover on this island (Battey *et al*. 2019), or a difference in responsiveness to warming between elevational generalists on species-poor tropical islands versus elevational specialists on species-rich mainlands (see above). Conversely, some temperate zone communities are closely tracking recent warming (Kelly & Goulden 2008; Menéndez *et al*. 2014).

### Absolute upslope shifts

We were motivated to test whether latitudinal position explains observed variation in temperature tracking. This approach standardizes species’ range shifts to the amount of local warming they have experienced. However, it is also true that some places on Earth are warming far faster than others, and that the overall rates of warming tend to be lower in the tropics (IPCC 2014). We therefore tested whether there were latitudinal differences in overall rates of upslope shift. For the tree plot dataset, we find that absolute responses to warming (thermophilization rates) are higher in the tropics, indicating that tropical tree communities are disproportionately responsive to warming regardless of metric. For range shift data, we find no significant difference between absolute responses to warming between latitudinal zones, but estimated shifts are slightly greater in the tropics (24.7 ± 6.33 m/decade) than the temperate zone (14.7 ± 2.88 m/decade). These values are similar to recent studies that have reported montane species are shifting upslope by 20.3 – 20.9 m/decade (Rumpf *et al*. 2019) and 17.8 m/decade upslope (Lenoir *et al*. 2020). However, these recent reports are substantially higher than the average overall shift for communities of 11.1 m per decade reported nearly a decade ago (Chen *et al*. 2011), which was itself double the estimate of 6.1 m per decade reported nearly two decades ago (Parmesan & Yohe 2003). Hence, as temperatures continue to warm and more datasets describing recent elevational range shifts are published (e.g. 93 communities in the present study versus 30 in the Chen et al. 2011 study), estimated rates of upslope shift continue to increase, likely reflecting an accelerating response of montane species to an accelerating driver of change.

### Conservation implications

The latitudinal gradient in temperature tracking we document has multiple implications for the conservation of montane floras and faunas, though we note that we found large variation within tropical and temperate regions. In the temperate zone, species’ upslope shifts are—on average— lagging far behind those expected given local warming. This indicates that acclimation and adaptation, rather than elevational shifts, may likely be key processes in determining how continued warming will lead to changes in population size for temperate montane species. It is an open question whether adaptation and acclimation will be able to keep pace with rates of warming that are unprecedented in recent evolutionary time (Visser 2008; Feeley *et al*. 2012). In contrast, the strong—on average—temperature tracking of tropical montane species indicates that continued warming is likely to lead to further upslope shifts, at least when protected elevational corridors provide suitable habitats at higher elevations. Upslope shifts’ consequences for populations will depend on the height and geometry of mountains. Upslope shifts may reduce population sizes when mountains are shaped like pyramids, with progressively less land at higher elevations, but could lead to population increases in cases where large areas of habitat exist in high elevation plateaus (Elsen & Tingley 2015). Most tropical mountains, however, lack high-elevation plateaus with suitable habitat, implying that upslope shifts will generally lead to progressive declines in population size that, unchecked, may ultimately lead to extirpations. That is, the “escalator to extinction” may run faster in the tropics. Such mountaintop extinctions are particularly likely for tropical species found only on single mountains or small mountain ranges that are of moderate height (Raxworthy *et al*. 2008). Notably, such mountaintop extinctions may occur well below the actual mountaintop, as pervasive anthropogenic modifications of high-elevation tropical systems effectively limit the ability of the tree line to shift upslope (Rehm & Feeley 2015). Indeed, local extinctions and range contractions associated with recent warming appear to be more common in tropical montane species than in temperate montane species (Wiens 2016; Freeman *et al*. 2018b).

The sixth mass extinction in Earth’s history is now underway (Ceballos *et al*. 2017). The tropics have the highest species diversity of any biome, and tropical mountains have the highest diversity of all (Rahbek *et al*. 2019). The relatively small temperature changes in the tropics should minimize the impact of climate change, but the disproportionate responsiveness of tropical montane species—on average—have instead potentially placed whole biotas on an escalator to extinction, though species-specific responses remain highly variable. The degree to which predictions of widespread local extirpations and species extinctions in tropical mountains (Şekercioĝlu *et al*. 2012) come true will depend on our ability to protect elevational corridors that enable species to persist while shifting upslope (Feeley & Rehm 2012) and, ultimately, on whether humanity is able to slow global warming.

## Acknowledgements

We thank the many scientists who have published range shift data and measured forest inventory plots. The Feeley, Schluter and Zhu labs, R. Yorque, J. Lembrechts and two anonymous reviewers provided useful comments that improved this manuscript. BGF was supported from postdoctoral fellowships from Banting Canada (379958), and the Biodiversity Research Centre, and YS was supported by a Regents’ fellowship from the University of California, Santa Cruz. KZ and KJF are supported by the US National Science Foundation (NSF grants DEB 1926438 to KZ and DEB-1350125 to KJF).

